# Geomosaic: a flexible bioinformatics platform integrating complementary metagenomic analyses from sequencing reads to genomes

**DOI:** 10.64898/2026.09.05.749574

**Authors:** Davide Corso, Edoardo Taccaliti, Bernardo Barosa, Donato Giovannelli

## Abstract

Metagenomic analyses can be performed at multiple analytical levels, including read-based, assembly-based, and genome-resolved approaches, each capturing complementary biological information while introducing distinct analytical biases and trade-offs. However, existing workflows are commonly optimized for a single analytical strategy, making it difficult to integrate these complementary representations within a unified, reproducible framework. Here we present Geomosaic, a modular framework that integrates complementary analytical representations of metagenomic data, from reads to genomes, within a single scalable, customizable, and reproducible workflow. Built on a graph-based architecture implemented in Snakemake, Geomosaic enables users to construct complete end-to-end workflows or execute individual analytical modules while selecting among interchangeable software packages. The framework supports read preprocessing, quality control, taxonomic and functional profiling, assembly, genome reconstruction, genome-resolved annotation, custom HMM-based analyses, and automated downstream result aggregation. Automatic generation of execution scripts, modular workflows, and multiple analysis entry points make Geomosaic accessible to researchers approaching metagenomic analyses for the first time, while providing the flexibility and control required by expert users. Native support for HPC environments enables efficient analysis of datasets ranging from individual projects to large-scale metagenomic surveys. Rather than treating read-, assembly-, and genome-resolved metagenomics as alternative analytical strategies, Geomosaic integrates them as complementary representations of the same biological system, allowing users to move seamlessly between community-wide patterns and organism-resolved functional interpretation. By combining workflow flexibility, computational reproducibility, standardized analysis-ready outputs, and extensive documentation, Geomosaic provides a unified platform for environmental metagenomic analyses and facilitates reproducible downstream ecological and evolutionary investigations.

**GRAPHICAL ABSTRACT:** Geomosaic integrates complementary read-, assembly-, and genome-resolved metagenomic analyses within a modular and scalable workflow. Its mosaic-like architecture combines flexible tool selection and HPC-ready execution, supporting metagenomic analyses from community-wide profiling to genome-resolved characterization.

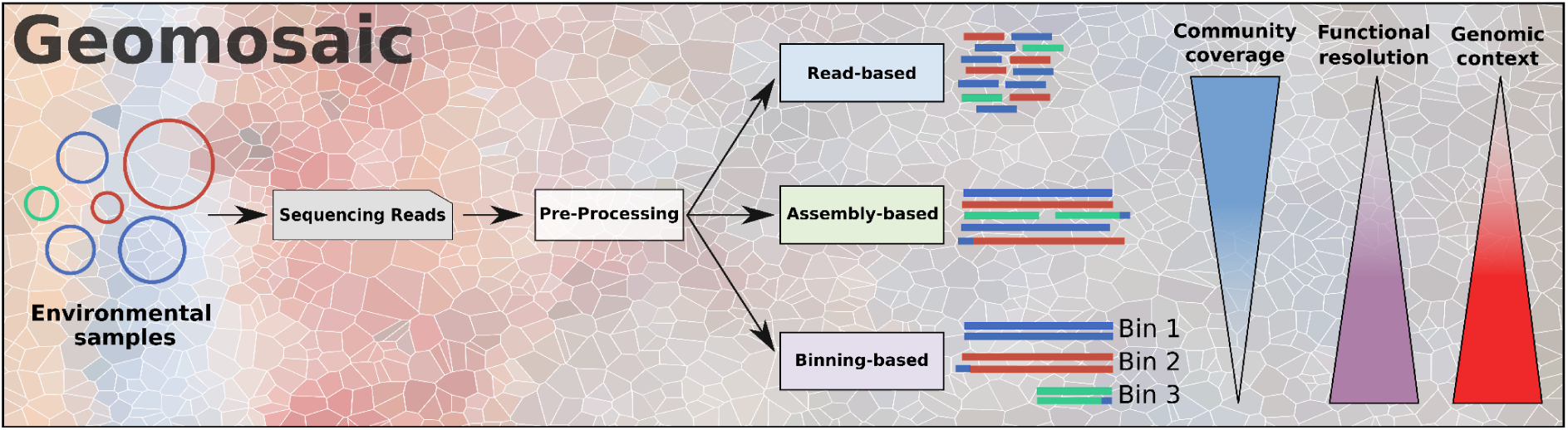

**HIGHLIGHTS:**

- Geomosaic integrates read-, assembly-, and genome-resolved metagenomic analyses in a single modular platform.
- A graph-based architecture enables flexible tool selection, modular execution, and extensible workflows.
- Automated workflow generation and HPC-ready scripts support both entry-level and expert users.
- Redox Metabolic and Metal Plasticity Indexes provide additional ecological interpretation of metagenomic functional potential.

## INTRODUCTION

Next-generation sequencing (NGS) technologies have revolutionized biological science research by enabling direct characterization of microbial communities without relying on cultivation. Metagenomic sequencing, in particular, provides access to the taxonomic composition and functional potential of complex communities and has substantially expanded our understanding of uncultured microbial diversity and its ecological roles (Lloyd et al., 2018; Parks et al., 2017; Riesenfeld et al., 2004).

The increasing volume of metagenomic data has, however, made analysis progressively more computationally demanding and methodologically complex. This trend is driven not only by the decreasing cost of sequencing and the resulting increase in sample numbers, but also by the growing use of computationally intensive strategies such as individual and co-assemblies, co-binning, cross-sample read-mapping, and the integration of newly generated and publicly available datasets within the same analysis (Li et al., 2015; Nurk et al., 2017). Modern metagenomic studies typically require multiple sequential and parallel steps, including read preprocessing, taxonomic and functional profiling, assembly, gene prediction, genome reconstruction, annotation, and abundance estimation. These analyses increasingly rely on High-Performance Clusters (HPC) infrastructures and require familiarity with bioinformatics tools, workflow management, software dependencies, and computational resource allocation (Ladoukakis et al., 2014; Uritskiy et al., 2018). Thus, getting insights from metagenomics data is a non-trivial task, as interdisciplinary knowledge, practical expertise, and multi-step analyses are required, either sequentially or in parallel (Ladoukakis et al., 2014).

Importantly, metagenomic data can be interrogated through different analytical representations that retain complementary biological information and are affected by different recovery biases. Read-based analyses preserve the broadest representation of the sequenced community (Goussarov et al., 2022) and are therefore well suited for semi-quantitative comparisons of taxonomic and functional patterns across samples, although they generally provide limited sequence completeness and genomic context. Assembly-based approaches reconstruct longer genomic fragments, increasing sequence completeness and enabling gene neighbourhood and contig-level analyses, but they describe only the fraction of the community that can be successfully assembled (Goussarov et al., 2022). Genome-resolved metagenomics further associates genes and functions with reconstructed microbial populations through metagenome-assembled genomes (MAGs), providing organism-level genomic context while introducing additional biases associated with assembly, binning, dereplication, and genome-quality filtering. Consequently, assemblies and MAGs should be interpreted as progressively selected subsets of the original sequenced community rather than as more accurate representations of it (Nelson et al., 2020). This distinction is particularly important for community-level metrics: for example, diversity estimates calculated from MAG collections reflect the diversity of the recovered genomic fraction rather than the entire microbial community. Because genome recovery depends on assembly and binning performance, which itself is influenced by community complexity and sequencing characteristics, MAG-based diversity estimates should not necessarily be interpreted as direct estimates of community diversity (Meziti et al., 2021; Nelson et al., 2020). Consequently, biological inferences derived from reads, assemblies and MAGs should be interpreted in the context of the biological entity they represent: the sequenced community, the assembled fraction of that community, and the recovered genomes, respectively. These approaches therefore represent complementary analytical layers that address different biological questions, rather than alternative levels of analytical quality.

Several metagenomic pipelines have been developed to automate subsets of these analyses (Ladoukakis et al., 2014), including read-based functional profiling, assembly and annotation workflows, and genome-resolved reconstruction (Garfias-Gallegos et al., 2022; Tadrent et al., 2023; Tamames & Puente-Sánchez, 2019; Yepes-García & Falquet, 2024). However, existing workflows are frequently optimized around a specific analytical representation or predefined sequence of tools. Moving between read-, assembly-, and genome-resolved analyses, combining their outputs, or replacing individual tools can therefore require separate pipelines, additional scripting, or substantial reconfiguration.

Here we present Geomosaic, a customizable metagenomic pipeline, designed to be automated, modular, and scalable, both on data computation and tool integration, thus reducing the complexity and providing a user-friendly approach to metagenomics. Geomosaic was designed to accommodate both users approaching metagenomic analysis for the first time and more experienced users requiring greater control over workflow design. Ready-to-run scripts, modular execution, and multiple entry points reduce the amount of manual configuration required, while preserving the flexibility to modify parameters, execute individual modules, or construct complete workflows. It is designed to be executed directly on HPC clusters and provides ready-to-run scripts using SLURM (Yoo et al., 2003) or GNU Parallel (Tange, 2023) technologies. In addition, its mosaic-like framework allows expert users to extend the pipeline by adding new modules or integrating alternative tools using Snakemake (Mölder et al., 2021), providing a simple route for the community to contribute to the future development of Geomosaic.

Beyond workflow execution, Geomosaic is open-source, available on GitHub (https://github.com/giovannellilab/Geomosaic) under the GPLv3 license and designed to produce standardized, analysis-ready outputs that facilitate ecological, evolutionary, and comparative interpretation across samples and analytical scales. A dedicated gathering layer integrates results generated across samples into structured matrices and tables suitable for downstream statistical analysis and visualization. These outputs can be explored using an accompanying Geomosaic Cookbook containing reproducible R and Python workflows for data exploration, visualization, and figure generation (https://github.com/giovannellilab/geomosaic_cookbook). Geomosaic therefore provides a unified framework for moving from raw metagenomic sequences to complementary community-, contig-, and genome-resolved biological information while preserving flexibility in analytical design.

## RESULTS

Geomosaic is a standalone pipeline that can be installed as a Conda environment, requiring no additional manual configuration, as shown in Table 2 together with its main features. It is designed to provide users with flexible control over the workflow, allowing customization of metagenomic analyses without imposing a fixed set of analytical steps. Users can choose to run a single module, execute the entire workflow from start to end, or combine both approaches depending on their needs.

Its modular structure enables the selection of a preferred software package for each analysis module, with the option to adjust, add, or modify parameters through configuration files within the *gm_user_parameters* folder. Once the pipeline has been customized, Geomosaic automatically assembles the selected modules and packages into a valid Snakemake workflow, removing incompatible modules and ensuring that dependencies are respected. Moreover, a core technical strength of Geomosaic, enabled by the integration of Snakemake, lies in its ability to use independent software packages for each module. By defining and isolating specific Conda environments for each tool during the initial configuration, the pipeline minimizes dependency and version conflicts.

Geomosaic supports extensive sample-level parallelization and is optimized for High-Performance Computing (HPC) environments. Each sample is assigned a dedicated working directory, simplifying the monitoring of computations and the organization of intermediate and final outputs. Parallel execution can be achieved using either GNU Parallel or automatically generated SLURM scripts, providing ready-to-run job submission files compatible with different computational infrastructures.

The pipeline accommodates novice users, who can execute workflows without modifying the underlying code, as well as advanced users, who can further adapt scripts and parameters as needed. The technical design of Geomosaic also facilitates community contributions, as new tools can be integrated by adding Snakemake rules and corresponding configuration entries using the provided documentation.

### Geomosaic commands

Geomosaic provides a set of user-friendly commands that guide the execution of the pipeline in a modular and reproducible way. Each command includes a built-in help section, offering clear instructions and configuration options. The workflow is organized around four core commands: *setup*, *workflow*, *unit*, and *prerun*, which are used to prepare, configure, and execute the analysis. In addition, an optional *gather* command allows users to merge results across samples into unified tables for downstream analyses. Together, these commands provide both simplicity for novice users and fine-grained control for advanced users, while maintaining flexibility, reproducibility, and integration with High-Performance Computing environments.

#### Geomosaic setup

The initial setup of Geomosaic organizes the working directory and prepares all provided sequencing samples for downstream analysis. The command takes as input a directory containing raw paired-end reads and a tabular sample table specifying the forward and reverse read files together with a sample identifier. The output of this step consists of a project-specific working directory containing a subdirectory for each unique sample and a setup file (e.g., *gmsetup.yaml*) that stores essential configuration information for reproducibility. This structure ensures consistent organization of reads across all subsequent pipeline steps and simplifies workflow execution across multiple samples. For example, executing the setup command on a directory of raw reads with a corresponding sample table results in individual sample folders, each containing the corresponding R1 and R2 files ready for analysis.

#### Geomosaic workflow

The *workflow* command in Geomosaic allows users to interactively select the analysis modules and corresponding software packages to create a customized pipeline. During this step, users are prompted to choose the preferred tool for each module; the list of available modules is dynamically updated depending on any modules that are skipped, ensuring that dependent modules that can no longer be executed are removed from the workflow. Once selections are finalized, Geomosaic generates a complete Snakemake workflow within the working directory, including a Snakefile containing the rules for the chosen modules, a *config.yaml* file with execution parameters, and an auxiliary *Snakefile_extdb.smk* for downloading and setting up external databases required by the selected tools. This approach provides a flexible and reproducible pipeline tailored to the user’s analytical choices while maintaining consistency across multiple samples or datasets.

#### Geomosaic unit

The *unit* command provides an alternative mode of execution in Geomosaic, allowing the user to run one module at a time rather than constructing the full workflow. This approach offers fine-grained control over the pipeline, enabling users to monitor the status of each job, inspect logs for errors, and selectively ignore failed samples in subsequent steps. For each execution of the *unit* command, Geomosaic generates a dedicated Snakefile (*Snakefile_unit.smk*) containing the rules for the selected module, a corresponding *config_unit.yaml* file for Snakemake execution, and, if required, a Snakefile for setting up external databases (*Snakefile_extdb.smk*).

By executing modules individually and reviewing logs between steps, users can inspect the outcome of each stage, adjust parameters, or rerun modules as needed without rerunning the entire workflow. This strategy is particularly useful for complex pipelines or large datasets, where intermediate failures may occur in specific samples, and provides a flexible framework to manage and control modular execution. For example, after running a module such as “*binning_derep”* with the unit command, a subsequent *prerun* step can be configured to exclude failed samples while specifying a dedicated log folder, maintaining reproducibility and transparency in the analysis.

#### Geomosaic prerun

The *prerun* command prepares Geomosaic for execution on high-performance computing clusters by automatically installing all required conda environments and generating ready-to-use execution scripts. The commands to execute these scripts are displayed on the screen in the correct order, allowing the user to submit jobs directly to the cluster (e.g., via *sbatch* for SLURM) without additional setup. Geomosaic accounts for dependencies such as external databases required by specific modules: if any chosen module needs pre-downloaded resources, the *prerun* command will show the correct order to fetch these external databases first, and then execute the main scripts for the analysis steps. Users can select SLURM or GNU Parallel for job submission, with optional specifications for memory, threads, partitions, or number of concurrent jobs. The *prerun* step can be applied after either a *unit* or a *workflow* command, ensuring that all modules and packages are correctly configured before execution. This design facilitates reproducibility, efficient resource management, and a fully guided execution of individual modules or complete workflows on a cluster.

#### Geomosaic gather

The *gather* command is an optional step designed to integrate results from multiple samples into unified, ready-to-use tables, for instance in a “samples × observations” format. This facilitates downstream analyses by providing pre-computed, merged datasets that summarize all relevant outputs. Not all modules generate gatherable results; only specific packages support this functionality. The modular design of Geomosaic allows users to selectively execute *gather* for the packages of interest, and it also enables contributions from the community to extend *gather* support to additional modules. This approach maintains flexibility while promoting streamlined data integration for downstream analyses and visualization.

#### Redox Metabolic and Metal Plasticity Indexes

Geomosaic introduces two novel metrics, the Redox Metabolic Index (RMI) and the Metal Plasticity Index (MPI), designed to quantify the functional potential and potential flexibility of microbial communities directly from read-based data. These indices are computed at the read-stream level to maximize the capture of latent metabolic diversity that is often reduced during assembly and binning. For any key gene identified via the fmh-funprofiler package, the module maps the corresponding KEGG Orthology (KO) identifier to a manually curated database that assigns electron donor or acceptor roles to specific biogeochemical substrates and their associated catalytic metal cofactors.

The module generates two outputs: a summary table reporting the calculated RMI and MPI values for each sample together with a concise inventory of the unique electron donors and acceptors identified, and an extended table that expands the results into a multi-row format, with each entry representing a specific metal–KO–substrate association. These structured outputs are formatted for direct integration into downstream visualization and analysis, or to correlate the indices with environmental variables and geochemical gradients.

Both indices were compared across the four sampled biomes (host-associated, seawater, soil, and subsurface; n = 6 samples per biome, total n = 24) using the Kruskal-Wallis test, followed by pairwise Dunn’s tests with Benjamini-Hochberg correction (Fig. 5). For the RMI (Fig. 5A), the Kruskal-Wallis test revealed a significant overall difference among biomes (χ² = 16.93, df = 3, p < 0.001). Subsurface samples showed the highest RMI values, followed by seawater, host-associated, and soil samples, which displayed the lowest values overall. Pairwise Dunn’s tests indicated that subsurface samples differed significantly from soil (p.adj < 0.001) and from host-associated samples (p.adj = 0.010), while all remaining pairwise comparisons, host-associated vs. seawater (p.adj = 0.337), host-associated vs. soil (p.adj = 0.337), seawater vs. soil (p.adj = 0.075), and seawater vs. subsurface (p.adj = 0.075), were not statistically significant after correction.

For the MPI (Fig. 5B), the Kruskal-Wallis test also indicated a significant overall difference among biomes (χ² = 14.15, df = 3, p = 0.0027). As for RMI, subsurface samples exhibited the highest MPI values, clearly separated from the other three biomes, which showed comparatively similar and overlapping distributions. Dunn’s post-hoc tests confirmed that subsurface samples differed significantly from host-associated (p.adj = 0.0057), soil (p.adj = 0.0048), and seawater samples (p.adj = 0.026), whereas no significant differences were detected among host-associated, seawater, and soil samples in any pairwise combination (all p.adj > 0.5).

Across both indices, subsurface samples consistently formed a distinct, higher-value cluster relative to the other three biomes, which instead displayed broadly overlapping distributions among themselves (Fig. 5A–B). In the case of MPI, higher values reflect a greater diversity of alternative metal-dependent redox configurations encoded within the community, potentially increasing the range of metabolic options available under varying trace-metal availability.

### Data analysis using Gemosaic

Geomosaic allows a customizable and flexible analysis of metagenomic samples. The user can choose across three stream levels (read-based, assembly-based, and genome-resolved), depending on the type of information required for the investigation, as well as on the questions asked and underlying hypotheses. If the user is interested in broad ecosystem-level analysis, then a read- or assembly-based approach may be preferable. On the other hand, if the user is interested in microbial biogeography, genome-resolved functional profiling, or pangenomics, then a genome-resolved approach should be considered.

Considering functional annotations, it is possible to choose among multiple packages and tools across the three main streams (Table 1). At the read-based level, mi-faser, fmh-funprofiler, and ARGs-OAP, with the possibility to include a custom database, were integrated. For annotating assembled contigs, users can predict ORFs using Prodigal and annotate them using different approaches; KOFAM-Scan, reCOGnizer, eggNOG-mapper, and HMM-based annotations were integrated. For MAGs, Geomosaic allows the choice among different tools and packages, including Bakta, HMM search, reCOGnizer, and KOFAM-Scan.

**Table 1.** Overview of the analysis modules and software packages implemented in the current release of Geomosaic. Modules are organized according to their role within the workflow, including workflow management, read-based, assembly-based, genome-resolved, and downstream analysis components.

| Stream-level | Modules | Packages | References |
| --- | --- | --- | --- |
| **Read based** | Pre-Processing | fastp | (Chen, 2023) |
|  |  | Trim Galore | (Krueger et al., 2023) |
|  |  | Trimmomatic | (Bolger et al., 2014) |
|  | Functional Annotation | mi-faser (GS-21-all) | (Zhu et al., 2018) |
|  |  | fmh-funprofiler | (Hera et al., 2024) |
|  |  | ARGs-OAP with custom DB | (Yang et al., 2016) |
|  | Taxonomic Annotation | Kaiju | (Menzel et al., 2016) |
|  |  | MetaPhlAn | (Blanco-Míguez et al., 2023) |
|  | Redox Metal Indexes | redox metal plasticity index | Custom implementation |
| **Assembly based** | Assembly | metaSPAdes | (Nurk et al., 2017) |
|  |  | MEGAHIT | (Li et al., 2015) |
|  | Assembly Quality Check | QUAST | (Gurevich et al., 2013) |
|  | Read Mapping | Bowtie2 | (Langmead et al., 2009) |
|  |  | Bowtie2<br>Output without unmapped reads | (Langmead et al., 2009) |
|  |  | BBMap | (Bushnell, 2014) |
|  |  | BBMap<br>Output without unmapped reads | (Bushnell, 2014) |
|  | Read Coverage | CoverM (contigs) | (Aroney et al., 2024) |
|  | Taxonomic Annotation | Kraken2 | (Wood et al., 2019) |
|  | ORF Prediction | Prodigal | (Hyatt et al., 2010) |
|  | Domain Annotation | reCOGnizer | (Sequeira et al., 2022) |
|  | HMM Annotation | HMM search | (Eddy, 2011) |
|  | Functional Annotation | eggNOG-mapper | (Cantalapiedra et al., 2021) |
|  |  | KOfam Scan | (Aramaki et al., 2020) |
|  | Assembly Redox Metal Indexes | Assembly redox metal plasticity index | Custom implementation |
| **Binning based** | Binning | Multi-Binners (MaxBin2, MetaBAT2, SemiBin2) | (Kang et al., 2019; Pan et al., 2022; Wu et al., 2016) |
|  | Binning De-Replication | DAS Tool | (Sieber et al., 2018) |
|  | Bins Quality Assessments | CheckM | (Parks et al., 2015) |
|  | MAGs Retrieval | MAGs retrieval | Custom implementation |
|  | MAGs Functional Annotation | DRAM (Temporary disabled) | (Shaffer et al., 2020) |
|  |  | Bakta | (Schwengers et al., 2021) |
|  | MAGs Taxonomic Annotation | GTDB-Tk | (Chaumeil et al., 2022) |
|  | MAGs ORF Prediction | Prodigal | (Hyatt et al., 2010) |
|  | MAGs Domain Annotation | reCOGnizer | (Sequeira et al., 2022) |
|  | MAGs ORF Annotation | KOfam Scan | (Aramaki et al., 2020) |
|  | MAGs Coverage | CoverM (genome) | (Aroney et al., 2024) |
|  | MAGs HMM annotation | HMM search | (Eddy, 2011) |
|  | MAGS Redox Metal Indexes | Mags redox metal plasticity index | Custom implementation |

**Table 2.**
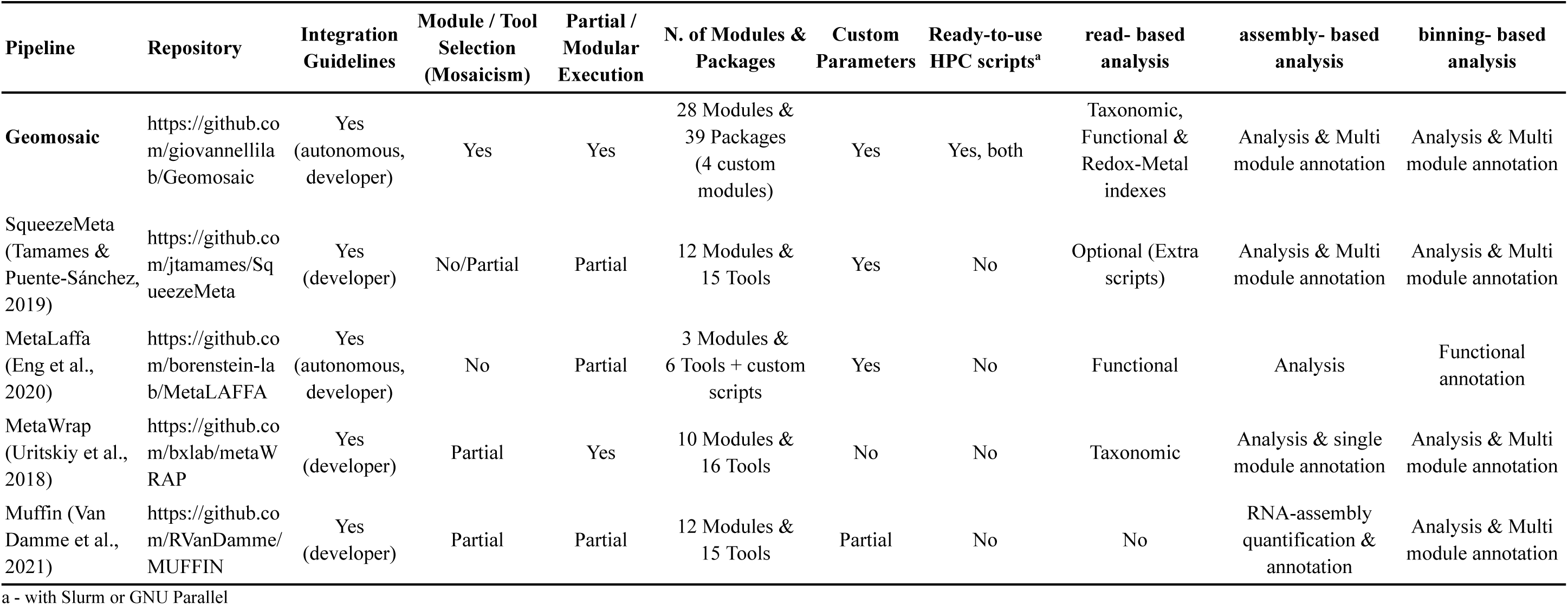
Comparison of Geomosaic with selected metagenomic analysis pipelines across workflow flexibility, analytical scope, HPC support, modularity, and extensibility.

To illustrate some possible analyses, we applied Geomosaic to samples from different biomes, including soil, seawater, host-associated, and subsurface environments (Supplementary Data X for sample information), using different modules. Within the read-based approach, we performed taxonomic (Fig. 3A) and functional profiling (Fig. 3C), using Kaiju and fmh-funprofiler, respectively. Furthermore, the newly integrated Redox Metabolic Index (RMI) and Metal Plasticity Index (MPI) analyses were performed. By applying these indexes across different biomes (Fig. 5), we observed that subsurface samples displayed higher RMI and MPI values.

**Figure 1.**
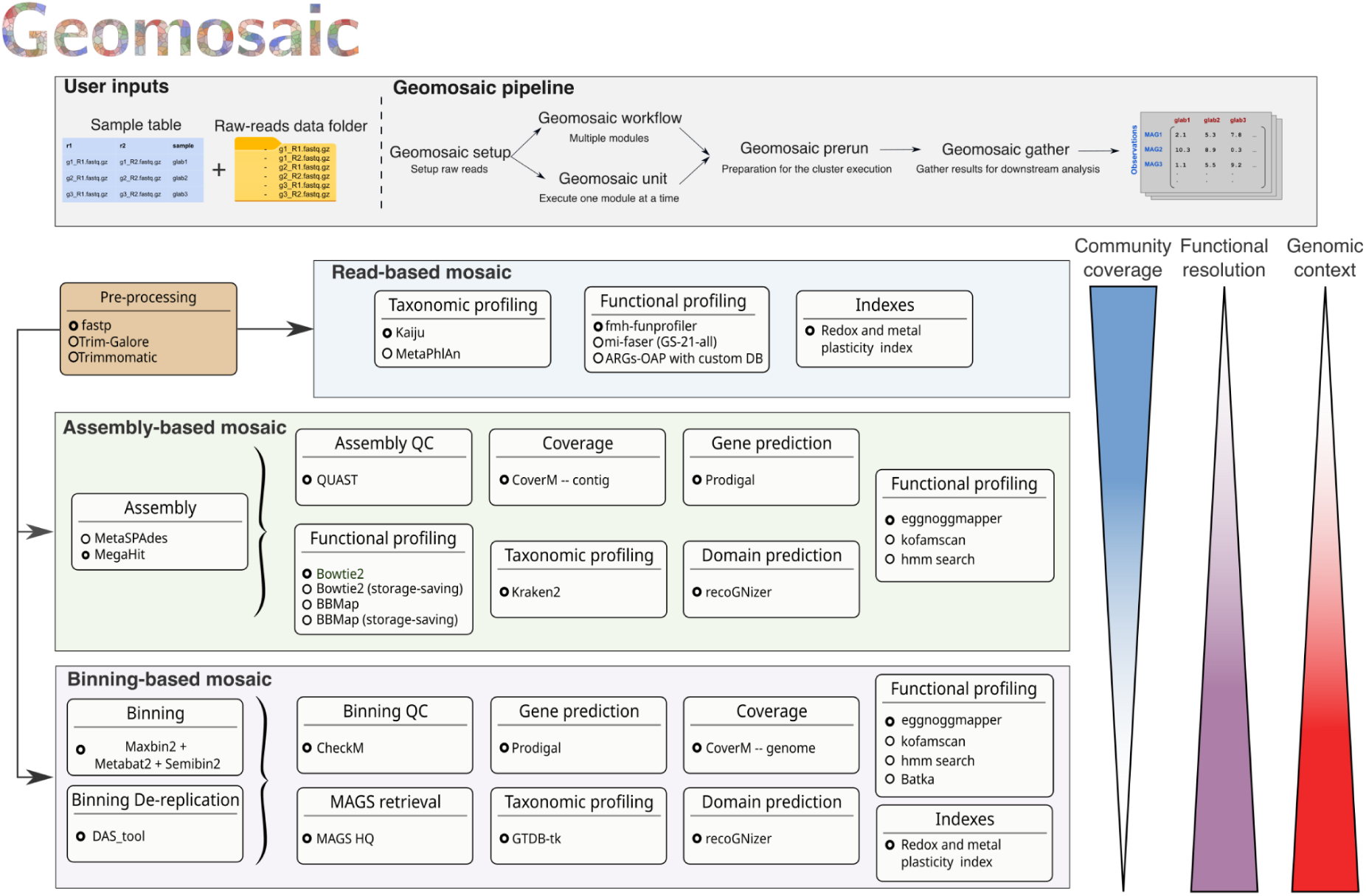
Overview of the Geomosaic workflow, highlighting the main user inputs, the three complementary analytical streams (read-based, assembly-based, and genome-resolved), and the corresponding outputs generated during pipeline execution.

**Figure 2.**
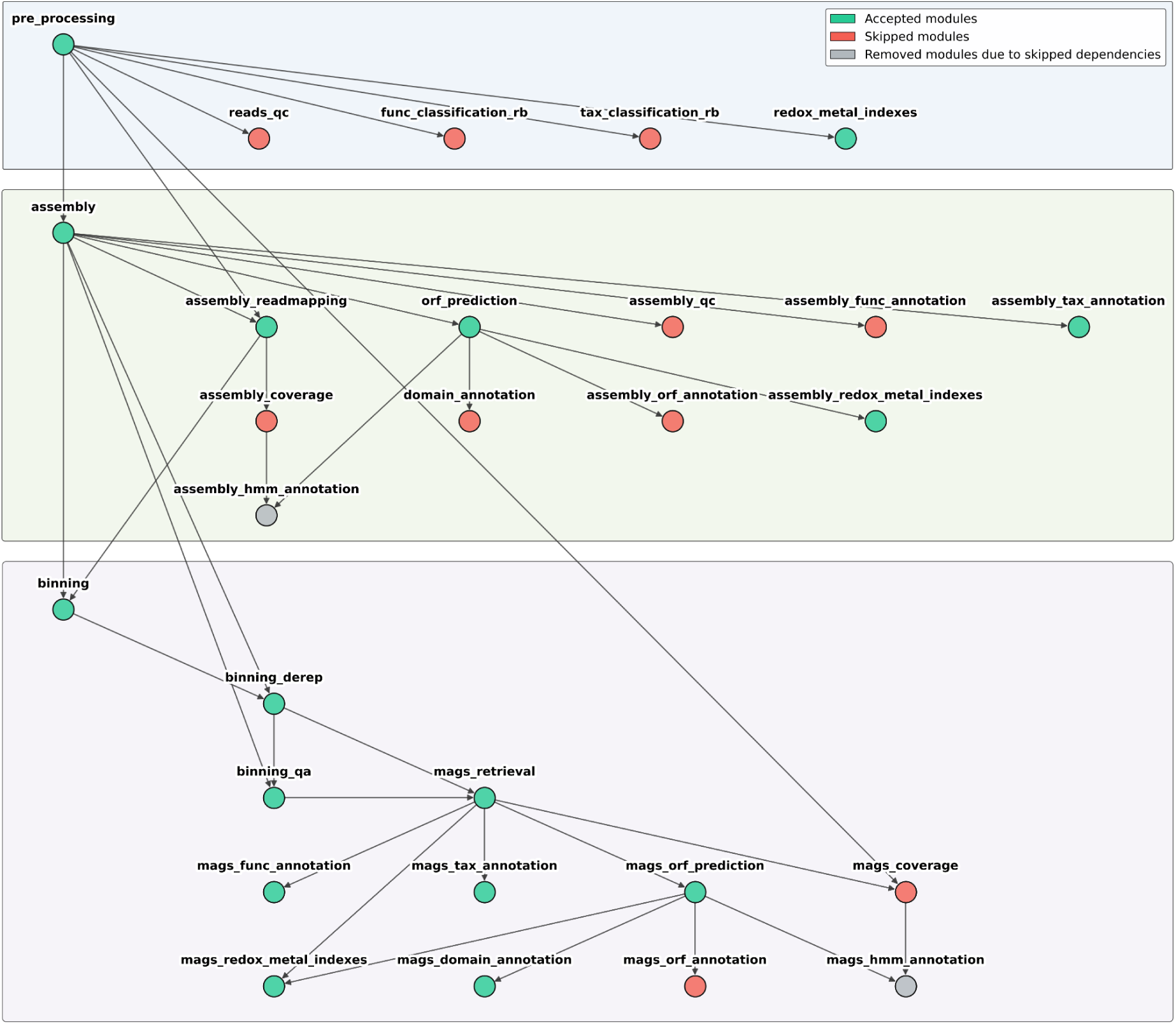
Example of a workflow graph generated by the *geomosaic workflow* command during the interactive selection of analysis modules. Each node represents a Geomosaic module and edges indicate dependencies between modules. Green nodes represent modules selected by the user, red nodes indicate modules explicitly skipped, and gray nodes correspond to modules automatically removed because one or more upstream dependencies were not selected. This dynamic graph-based approach ensures that only valid workflows are generated while providing immediate visual feedback on the consequences of each module selection.

**Figure 3.**
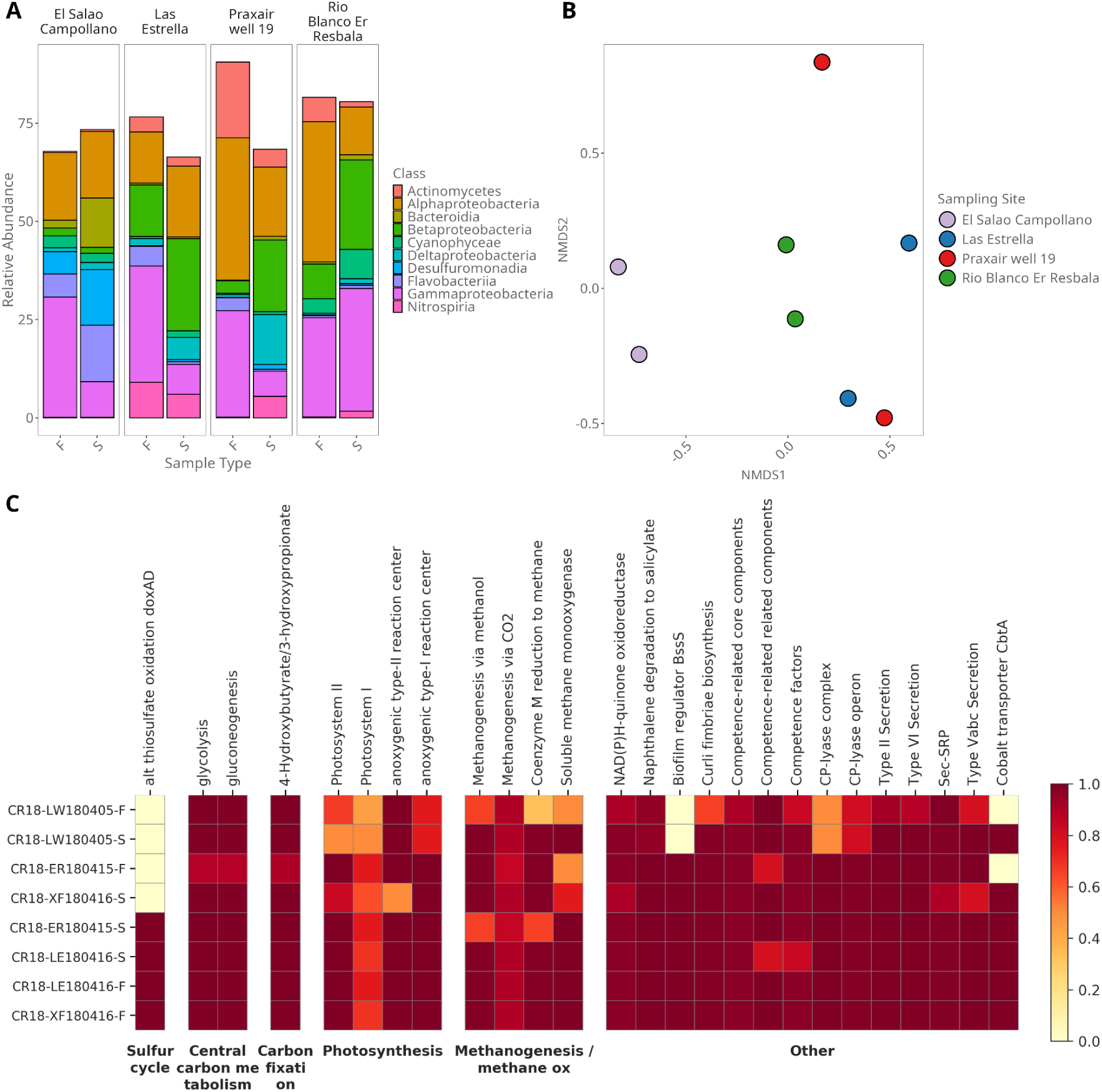
Example of downstream analyses and plots obtained using *gather* on the read-based analysis stream of Geomosaic and plotted using code from the Geomosaic Cookbook. A) Taxonomic distribution of sequencing reads obtained using the Kaiju module and visualized with the phyloseq R package. B) Principal Coordinate Analysis (PCoA) of taxonomic community composition, generated using the phyloseq R package. C) Distribution of a subset of functional genes, annotated using the KOFAM-Scan module and mapped to KEGG pathways using KEGG Mapper.

Taxonomic and functional annotation at the genome-resolved level was performed using GTDB-Tk (Fig. 4A) and the MAGs HMM annotation module to provide a comprehensive profiling of a selected sample (Fig. 4B).

**Figure 4.**
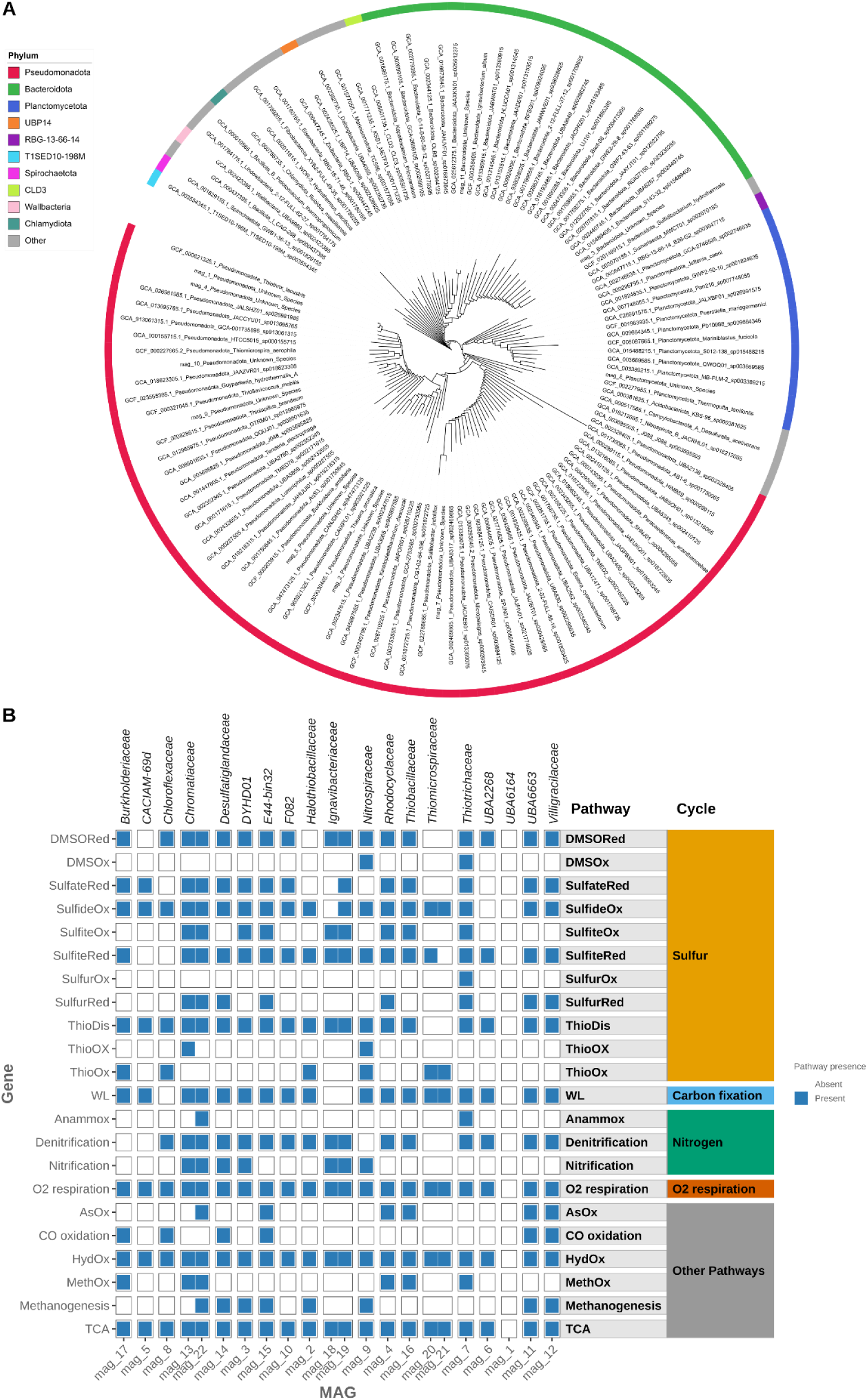
Examples of outputs obtained from the genome-resolved analysis stream of Geomosaic. A) Phylogenetic tree of the recovered MAGs, taxonomically classified using the GTDB-Tk module and visualized with iTOL. B) Heatmap showing the distribution of key biogeochemical genes identified using the MAGs HMM Annotation (MAG-HMMA) module and visualized in R.

**Figure 5.**
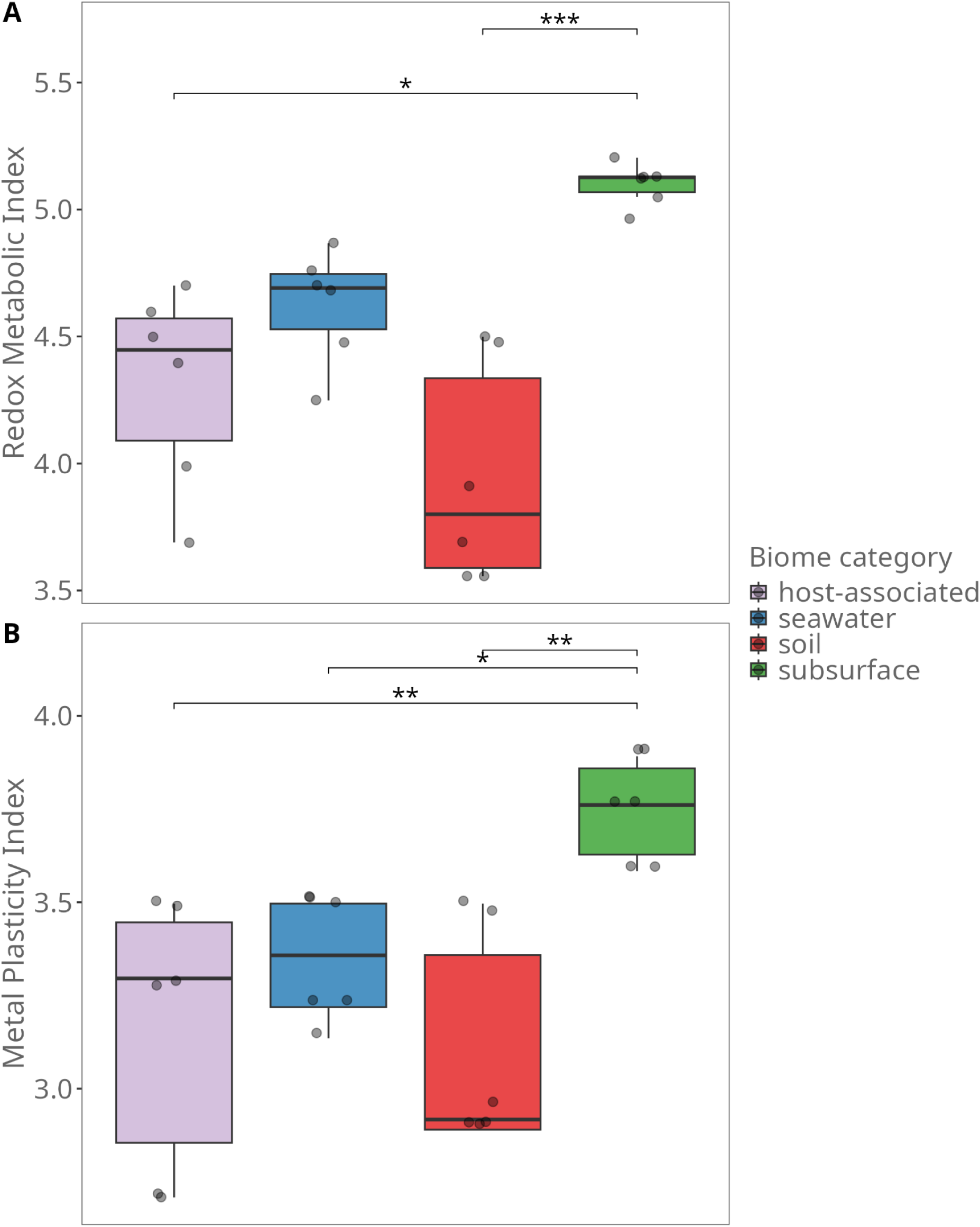
Variation in the Redox Metabolic Index (RMI) (A) and the Metal Plasticity Index (MPI) (B) across metagenomes from four distinct biomes (host-associated, seawater, soil, and subsurface), calculated using the read-based analysis stream of Geomosaic. Boxes represent the interquartile range, the central line indicates the median, whiskers extend to 1.5× the interquartile range, and individual points represent biological samples. Different asterisks indicate statistically significant differences according to Dunn’s post hoc test with Benjamini–Hochberg correction (*P* < 0.05).

### User documentation and tutorials

Users can access the complete Geomosaic documentation at https://giovannellilab.github.io/Geomosaic, which provides installation instructions, a step-by-step tutorial for setting up a first Geomosaic project, detailed command references, and practical guidelines for efficient workflow execution and reproducible analyses.

Moreover, a concise Geomosaic Cookbook for downstream analyses is available at https://github.com/GiovannelliLab/geomosaic_cookbook. The Cookbook provides documented Jupyter notebooks, Python and R scripts designed to operate directly on the outputs of the gather command, allowing users to explore, analyze, and visualize Geomosaic results and to reproduce the publication-ready figures presented in this work. Instructions and scripts for creating a dedicated Conda environment are also provided to facilitate reproducibility of the downstream workflows.

## DISCUSSION

Geomosaic was designed to integrate complementary representations of metagenomic data within a single scalable and customizable framework, while lowering the barrier to entry for users approaching metagenomic analysis. Rather than considering read-based, assembly-based, and genome-resolved metagenomics as competing analytical strategies, Geomosaic treats them as complementary representations of the same biological system, each retaining different dimensions of biological information. This was achieved in multiple ways: i) taking advantage of a modular graph structure that allows the user to choose among 28 different modules (Table 1), as well as integrating custom modules; ii) generating SLURM-ready scripts to run on HPC clusters; iii) enabling analysis across complementary analytical scales, from broad community-level read-based profiling to assembly-based and genome-resolved characterization; and iv) providing robust and comprehensive documentation detailing all commands and pipelines within Geomosaic.

Overall, and compared to similar tools (Table 2), Geomosaic distinguishes itself through its modularity and resource-management flexibility. While established tools often operate as rigid or partially flexible frameworks, Geomosaic is built on a graph-based architecture that allows the selection of specific software across 28 modules spanning three distinct stream levels of analysis (read-, assembly-, and genome-resolved), paired with straightforward tool parametrization. This structure allows users to combine multiple analytical representations within the same workflow while selecting the tools that best match their research questions.

This structure also enables the integration of new software components, providing a high degree of “mosaicism” that allows developers to extend the pipeline autonomously. This flexibility provides an advantage over tools with more constrained module integration. To address the bottleneck of computational scalability, Geomosaic was explicitly designed for HPC environments through integration of the Snakemake workflow management system. By providing ready-to-use, editable scripts for both shell-level task distribution via GNU Parallel and cluster-level resource management via SLURM, Geomosaic allows complex workflows to remain portable and efficiently executable across different computational infrastructures. This strategy allows researchers to scale analyses from local machines to high-performance clusters with minimal configuration while facilitating reproducibility and throughput.

An additional strength of Geomosaic is that it produces standardized, analysis-ready outputs that can be integrated across samples and analytical streams through the *gather* command. These outputs can be directly used for downstream ecological and comparative analyses and explored through the accompanying Geomosaic Cookbook, which provides reproducible R and Python workflows for data exploration, visualization, and figure generation. This reduces the gap between primary bioinformatic processing and biological interpretation, which can otherwise require substantial additional scripting and data reformatting.

In addition, Geomosaic includes dedicated modules for in-depth ecological interpretation at the read-based level, introducing two novel indexes, the Redox Metabolic Index (RMI) and the Metal Plasticity Index (MPI), as proxies for redox metabolic potential and potential flexibility in metal-dependent metabolic strategies within an environmental setting. These indexes are calculated by mapping enzymes representative of major metabolic pathways together with their corresponding metal cofactors. Previous studies have demonstrated the role of metal availability in constraining microbial communities and influencing the use of alternative respiratory pathways in both environmental and laboratory settings (Giovannelli, 2023; Ricciardelli et al., 2025). In our analyses, we found that both indexes were higher in subsurface samples and significantly different from several of the other investigated biomes. This could reflect the high diversity of electron donors and electron acceptors available in subsurface environments, together with the influence of water-rock reactions on trace-metal availability (Rogers et al., 2023; Tivey, 2007). In particular, higher MPI values indicate a greater diversity of alternative metal-dependent redox configurations encoded within the community, potentially providing a broader range of metabolic options under changing trace-metal availability. The higher RMI and MPI values may therefore reflect the distinct environmental niches present in such systems. We propose that these indexes can provide an additional layer of information for researchers aiming to characterize environmental metagenomes in greater detail.

Looking ahead, Geomosaic development will focus on major features such as the ability to import data at different stages (e.g., assemblies or co-assemblies) and the integration of modules for long-read metagenomic analysis. A further planned feature is a geomosaic plot command to generate charts and reports directly from the pipeline’s gathered files, extending the current downstream functionality provided through the Geomosaic Cookbook. To further increase reproducibility and portability across HPC architectures, Geomosaic will be encapsulated in a containerization system such as Docker or Singularity. Finally, an extensive geochemical pipeline will be integrated to provide dedicated scripts for processing geochemical data alongside the biological modules.

Taken together, these results show that Geomosaic offers a modular, scalable, and extensible framework that lowers the technical barrier to metagenomic analysis without sacrificing methodological flexibility. By combining complementary read-, assembly-, and genome-resolved analytical streams with graph-based module selection, HPC integration, standardized analysis-ready outputs, and custom ecological modules such as RMI and MPI, Geomosaic provides researchers with a tool that can adapt to the scale and specificity of their questions, from broad community-level screening to genome-resolved characterization. As the tool continues to evolve, incorporating long-read support, co-assembly import, containerization, enhanced downstream visualization, and geochemical data integration, Geomosaic is positioned to become a versatile resource for the broader community studying environmental microbiomes and biosphere-geosphere interactions.

## CONCLUSION

Geomosaic provides a flexible and extensible platform integrating complementary metagenomic analyses from reads to genomes. Its modular architecture and HPC-ready execution lower the technical barrier to metagenomic analysis while retaining flexibility for expert users. Together with standardized outputs and dedicated ecological indexes, Geomosaic provides a scalable framework for investigating microbial communities across diverse environments.

## MATERIALS AND METHODS

### Core graph structure

Geomosaic’s core structure is built upon a graph-based architecture which provides different features and advantages in the management of the integrated modules. In addition, it also represents a conceptual map that can be followed by the user during the creation of the workflow. In Geomosaic, each node of the graph represents an analysis module, while edges among modules describe their dependencies, thus providing intrinsic advantages in the construction of pipelines that require the handling of sequential input and output steps. Another major advantage of this architecture is its complementarity with Snakemake, which performs internal checks on the integrity of the pipeline based on the specified inputs and outputs.

By considering each node as an analysis module, Geomosaic can be integrated with an arbitrary number of packages defined by the user. These tools are considered independent alternatives within each module. As a consequence, the graph structure of Geomosaic allows the user to choose the package of interest and, based on these choices, creates the corresponding workflow. Specifically, by using the core graph, a queue data structure, and a module order established on their dependencies, Geomosaic provides a module selection that dynamically changes based on each choice. If the user decides to ignore a step, all dependent modules that can no longer be executed are removed from the queue and are not provided to the user.

The choice of a graph-based design was also made to lower the user barrier by tailoring the workflow to the analyses of interest. The current integrated modules in the graph are extensively explained in the Geomosaic documentation, which facilitates the understanding of step dependencies and thus the customization of the pipeline. Once the procedure is finished, the resulting workflow is a subgraph of the original structure, containing only the nodes representing modules chosen by the user.

Moreover, the modular structure of each analysis allows the integration of novel modules by defining their dependencies and packages, integrating the execution code using Snakemake syntax, and, when required, specifying external databases. We are aware that contributing to a third-party pipeline is not a trivial task; therefore, Geomosaic was designed to accommodate both users seeking a customizable workflow and users with bioinformatic expertise who wish to extend the framework. Full documentation on how to contribute novel modules is provided in the Geomosaic documentation.

### Implementation

Geomosaic was developed as a Python package, as Python provides libraries to handle command-line options and parameters, workflow construction based on the selected modules, graph data structures, and integration with Snakemake.

Snakemake is the framework used for workflow management of the modules included in the resulting pipeline. It provides a user-friendly and extensively documented interface, with a human-readable syntax based on rules, and supports reproducible and scalable data analyses. Moreover, for each rule, Snakemake allows the specification of an external Conda environment, typically defined through a configuration file, helping to avoid conflicts among dependencies required by different modules. For this purpose, for every integrated package we defined a corresponding Conda configuration file. In this way, when a rule is executed, the corresponding environment is used independently from those of other modules.

As described above, Geomosaic integrates different metagenomic analysis modules that can require substantial computational resources and was therefore designed to be executed on High-Performance Computing clusters. For this reason, we provide a command to create SLURM or GNU Parallel scripts for parallel execution of Geomosaic workflows across samples. Specifically, these scripts execute the workflow for each sample in parallel.

If SLURM is available on the HPC system, we suggest using the corresponding scripts, as SLURM provides scalable cluster management and job scheduling. It also allows users to specify computational resources such as execution time, CPUs, and memory. Since HPC systems may require additional SLURM directives, such as account or partition information, the generated SLURM scripts are intended to be editable by users to accommodate the requirements of their specific HPC environment. If SLURM is not available, users can instead generate scripts based on GNU Parallel. In this case, users must manually define the number of parallel jobs and the number of CPUs assigned to each computation.

### Collected Modules

We outlined three complementary streams of metagenomic analysis, which users can select and combine depending on their analytical goals, choosing among the available analysis modules and the integrated tools within each stream. In the read-based stream, annotations are performed directly on filtered reads, retaining the broadest representation of the sequenced community. In the assembly-based stream, analyses focus on the contigs reconstructed from those reads, providing increased sequence completeness and local genomic context. Finally, in the genome-resolved stream, functional and taxonomic characterization is carried out on the high-quality metagenome-assembled genomes (MAGs) recovered from the assemblies, providing organism-level genomic context. Using the modular core of Geomosaic, users can tailor the workflow based on their interests, taking into consideration the complementary information and trade-offs associated with each analytical stream.

Geomosaic was designed to optimize metagenomic analysis across these complementary levels. As already described, existing metagenomic workflows often focus predominantly on one type of output, particularly read-based profiling or genome-resolved analyses. However, there is a progressive reduction in the fraction of the original metagenomic signal retained through the sequential steps required to move from reads to assemblies and MAGs. Despite the importance of this workflow for reconstructing genomic context, we consider all data representations, including the raw and filtered reads, to contain potentially valuable and complementary information. Read-based annotations, assembly-based analyses, and genome-resolved results therefore address different biological questions and should not be considered alternative levels of analytical quality. For this purpose, we integrated different analysis modules, including multiple types of annotation, across all three stream levels: read-based, assembly-based, and genome-resolved.

All the collected programs were chosen mainly based on their installation method and compatibility with the Geomosaic framework. Specifically, Conda-based or PyPI-based installations were preferred because they can be readily integrated into the management of corresponding Conda environments by Snakemake. This facilitates reproducibility of the workflow and reduces dependency conflicts between modules. Currently, the graph structure is made up of 28 modules, integrating a total of 39 packages and custom implementations. All currently integrated modules are shown in Table 1.

### Custom modules developed for Geomosaic

#### Assembly HMM Annotation

An important feature of Geomosaic is the ability of the user to use their own annotation sources, and for that reason we enable the use of a user-compiled collection of Hidden Markov Models (HMMs).

This dedicated module to annotate open reading frames (ORFs) has been implemented in Geomosaic. Specifically, this module has been designed to run at the assembly-stream level. The ORF protein sequences are screened against a user-provided collection of HMM profiles using *hmmsearch* from the HMMER tool. HMMER searches are performed with the following parameters: *-o*, *--cpu*, and *--notextw*.

Results are then parsed using Biopython (≥1.82) to extract alignment strings, from which sequence similarity metrics are computed based on both identical and conserved matches reported in the HMMER alignment output as follows (Corso et al., 2026):

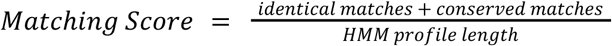

Conserved matches are defined as residues marked with a “+” in the HMMER alignment string, indicating conservative substitutions with positive log-odds emission scores expected by the profile, thereby capturing both strict sequence identity and biologically meaningful substitutions when assessing similarity between sequences and HMM models. To facilitate downstream quantitative analyses, annotation results are subsequently integrated with the ORF–contig mapping generated during gene prediction and with coverage estimates derived from read mapping. Coverage values obtained through multiple methods can be incorporated and are merged with the annotation results through contig identifiers. The final output consists of a comprehensive table reporting detected HMM models, associated ORF IDs, detailed alignment metrics (e.g., number of identical and conserved matches, corresponding similarity percentages, bitscore and E-values), contig identifiers, and contig-specific coverage values.

#### MAGs HMM Annotation

An analogous procedure was implemented to perform HMM-based annotation on ORFs predicted from MAGs. In this case, ORF protein sequences derived from each MAG are screened against the user-provided HMM profile collection using the same *hmmsearch* strategy from the HMMER tool, following the same parameters and parsing procedure described above. Similarity metrics based on identical and conserved matches are computed from the alignment output as previously described.

The resulting annotations are then integrated with MAG identifiers and coverage estimates for each sample, generating a table reporting HMM models, associated ORF IDs, detailed alignment metrics, MAG and contig identifiers, and MAG-specific coverage values.

#### Redox Metabolic Index

To quantify the overall redox metabolic potential within a given sample, we developed the Redox Metabolic Index (RMI). This metric is designed to capture the theoretical maximum number of possible combinations between electron donor and electron acceptor species identified in the metagenomic dataset.

For any key gene encoding a catalytic enzyme associated with a specific KEGG Orthology (KO) identifier, we define its metabolic role based on established literature as an acceptor and/or donor of electrons for specific biogeochemical substrates, as well as its corresponding catalytic metal cofactor, where applicable. For this purpose, a specifically designed curated database of biogeochemically relevant KOs was manually built to provide a further layer on top of the functional information available in the KEGG database. In particular, we define each KO as either a source of electrons or an electron acceptor or both for a known biogeochemical substrate on which it operates. We decided to also count non-catalytic components of the protein machinery as contributors to the index computation, since their detection in the metagenomic sample can provide additional evidence for the presence of the corresponding metabolic machinery.

We implemented this index specifically at the read-based stream level to retain the broadest representation of functional information, which is progressively reduced during subsequent assembly and binning processes. Functional profiling of the reads is performed using fmh-funprofiler to achieve rapid identification of metabolic potential.

The mathematical derivation of the RMI is rooted in Shannon entropy. For a community where the abundances of metabolic species are unknown, we first define the general entropy function:

[uamth2]

When no prior information is available, entropy is maximized by assuming an equal probability for each species 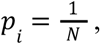 which simplifies the derivation to:

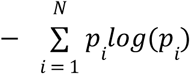

Consequently, the RMI is defined as the logarithm of the product of the number of available donor and acceptor substrate species, or equivalently as the sum of their individual logarithms:

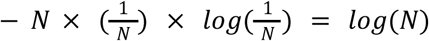

In this context, the donor (*N_d_*) and acceptor (*N_a_*) terms represent the number of unique donor and acceptor substrate species identified by mapping KEGG Orthology (KO) hits from the read-based functional annotation to the manually curated custom database encompassing the broader biogeochemical pathway landscape. The RMI and MPI (see below) can be calculated at both the whole-metagenome level (using either read- or assembly-based KO functional annotations) and the individual MAG level, enabling the same ecological framework to be applied across analytical scales, from microbial communities to reconstructed genomes.

#### Metal Plasticity Index

The Metal Plasticity Index (MPI) is built upon the RMI by incorporating an additional layer of information on the metal requirements at the catalytic cores of the identified donor- and acceptor-associated enzymes. Because metalloenzymes mediate critical electron-transfer reactions (Giovannelli 2023; Hay Mele et al 2023), the diversity of metal cofactors associated with alternative redox pathways can provide information on the potential of a community to respond to variations in trace-metal availability. We define the MPI as the logarithm of the number of unique, unordered metal pairs available for potentially coupled redox reactions:

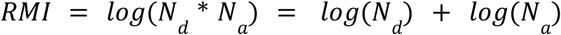

This index highlights the potential trade-off between metabolic specialization and flexibility. For instance, a system encoding only an Fe–Cu donor–acceptor combination relies on that specific metal configuration. Conversely, a system encoding several alternative redox combinations, such as Fe–Fe, Fe–Cu, and Fe–Mo, has a greater potential to maintain alternative electron-transfer pathways under changing trace-metal availability. This is consistent with experimental evidence showing that trace-metal availability can control the utilization of alternative terminal electron acceptors by microorganisms (Ricciardelli et al., 2025). The presence of metal-binding promiscuous proteins (Hay Mele et al 2023), which can incorporate alternative metals in place of the native cofactor, may further contribute to this potential flexibility.

To calculate the MPI, Geomosaic utilizes the separate lists of donor and acceptor species generated during the previous RMI calculation and identifies all unique connections between the metals associated with those specific substrates.

## AUTHOR CONTRIBUTIONS

**DC**: Conceptualization, Software, Methodology, Formal analysis, Writing – original draft, Visualization. **ET**: Software, Writing – review & editing. **BB**: Conceptualization, Writing – review & editing. **DG**: Conceptualization, Methodology, Supervision, Funding acquisition, Project administration, Writing – review & editing.

## ACKNOWLEDGMENTS

This work was supported by the European Research Council (ERC) under the European Union’s Horizon 2020 Research and Innovation Programme through the COEVOLVE project (Grant Agreement No. 948972) awarded to D.G. The authors gratefully acknowledge all members of the Giovannelli Lab who contributed to the testing, debugging, and continuous improvement of Geomosaic throughout its development. Their feedback, testing across diverse datasets, and suggestions were instrumental in refining the pipeline and improving its robustness and usability. The authors also thank Dr. Nathaniel Virgo for his insightful discussions and contributions to the mathematical formulation of the Redox Metabolic Index (RMI) and the Metal Plasticity Index (MPI), and Guillermo Climent Gargallo for integrating the Bakta annotation tool into Geomosaic.

## ETHICS STATEMENT

No animals or humans were involved in this study.

## DATA AVAILABILITY STATEMENT

Geomosaic is an open-source software package released under the GNU General Public License v3.0 (GPLv3). The source code, documentation, installation instructions, and example workflows are publicly available through the GitHub repository: https://github.com/giovannellilab/Geomosaic. Versioned software releases are permanently archived on Zenodo and assigned a DOI to ensure long-term reproducibility. The current version is v1.5.2 available with DOI https://doi.org/10.5281/zenodo.22307030 and can be directly installed using Conda. Comprehensive user documentation is available at https://giovannellilab.github.io/Geomosaic, and reproducible downstream analysis workflows are provided through the Geomosaic Cookbook at https://github.com/giovannellilab/geomosaic_cookbook. The metagenomic datasets used in this study are publicly available through the NCBI Sequence Read Archive and the European Nucleotide Archive. Subsurface samples were previously described and published by Paul et al., (2023) and deposited under BioProject PRJNA914269, which is part of the larger umbrella project PRJEB55081. Complete accession information — including run, study, and sample accessions — for all samples analyzed in this study is provided in Supplementary Table S1 available at https://doi.org/10.5281/zenodo.22339508.

## SUPPORTING INFORMATION

**Supplementary Table S1.** Metagenomic datasets used for demonstration and benchmarking of Geomosaic, provided as *Table_S1_metagenomic_datasets.csv*. The table lists the 24 publicly available datasets analyzed in this study, spanning four biomes (host-associated, seawater, soil, and subsurface), and reports for each the run accession, biome/metagenome type, study accession, and sample accession. Available at https://doi.org/10.5281/zenodo.22339508.

**Supplementary Table S2.** Curated KO–substrate–redox role–metal cofactor database, provided as *Table_S2_KO_substrate_redox_metal_database.csv*. This database links KEGG Orthology (KO) identifiers to their associated substrates, redox roles, and metal cofactors, and was used as the reference set for calculating the Redox Metabolic Index (RMI) and Metal Plasticity Index (MPI). Available at https://doi.org/10.5281/zenodo.22339508.

## COVER

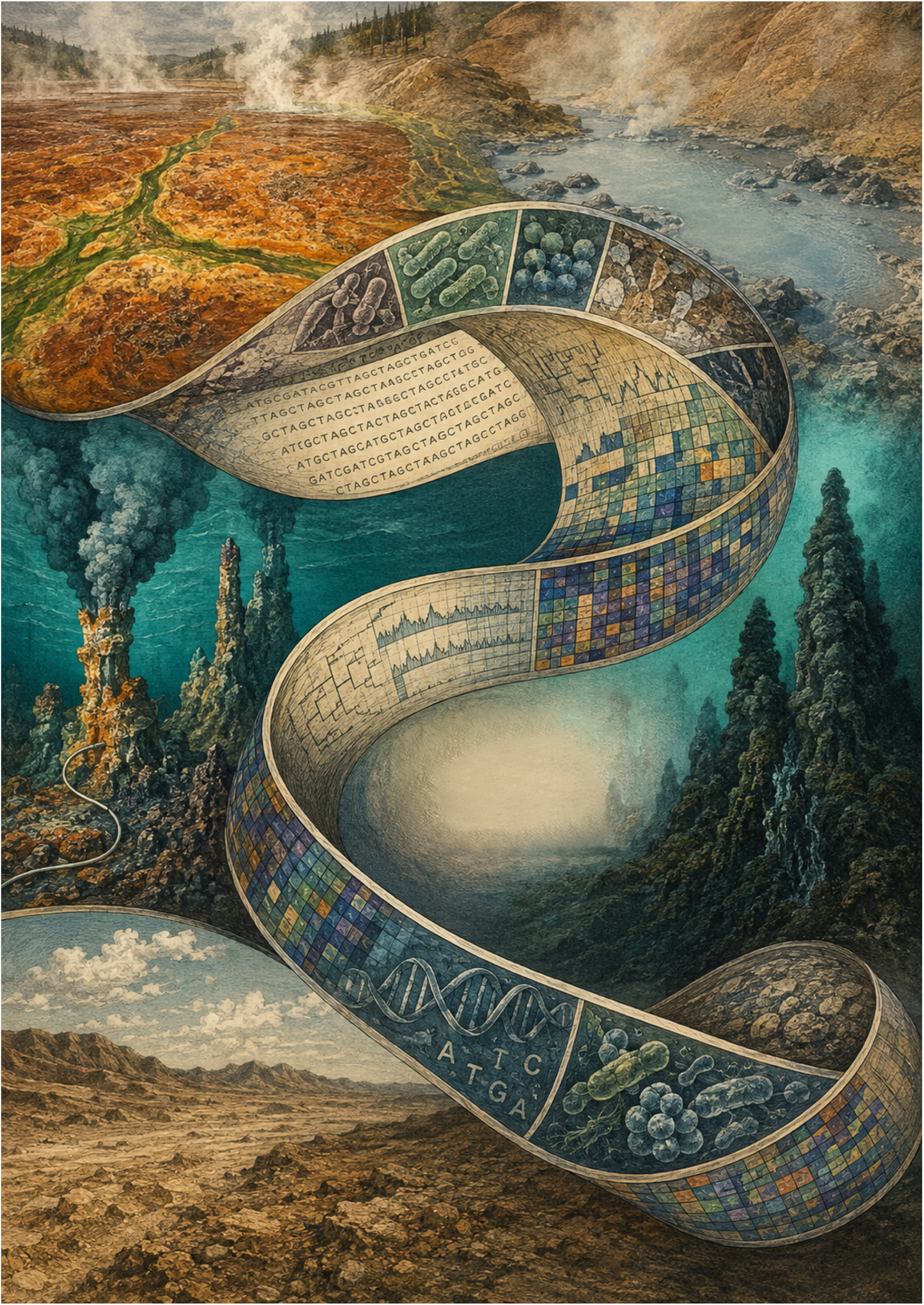

## REFERENCES

Aramaki, T., Blanc-Mathieu, R., Endo, H., Ohkubo, K., Kanehisa, M., Goto, S., & Ogata, H. (2020). KofamKOALA: KEGG Ortholog assignment based on profile HMM and adaptive score threshold. Bioinformatics, 36(7), 2251–2252. 10.1093/bioinformatics/btz859

Aroney, S. T. N., Newell, R. J. P., Nissen, J., Camargo, A. P., Tyson, G. W., & Woodcroft, B. J. (2024). *CoverM: Read coverage calculator for metagenomics* (Version v0.7.0) [Computer software]. [object Object]. 10.5281/ZENODO.10531253

Blanco-Míguez, A., Beghini, F., Cumbo, F., McIver, L. J., Thompson, K. N., Zolfo, M., Manghi, P., Dubois, L., Huang, K. D., Thomas, A. M., Nickols, W. A., Piccinno, G., Piperni, E., Punčochář, M., Valles-Colomer, M., Tett, A., Giordano, F., Davies, R., Wolf, J., … Segata, N. (2023). Extending and improving metagenomic taxonomic profiling with uncharacterized species using MetaPhlAn 4. Nature Biotechnology, 41(11), 1633–1644. 10.1038/s41587-023-01688-w

Bolger, A. M., Lohse, M., & Usadel, B. (2014). Trimmomatic: A flexible trimmer for Illumina sequence data. Bioinformatics, 30(15), 2114–2120. 10.1093/bioinformatics/btu170

Bushnell, B. (2014). BBMap: A Fast, Accurate, Splice-Aware Aligner. https://www.osti.gov/biblio/1241166

Cantalapiedra, C. P., Hernández-Plaza, A., Letunic, I., Bork, P., & Huerta-Cepas, J. (2021). eggNOG-mapper v2: Functional Annotation, Orthology Assignments, and Domain Prediction at the Metagenomic Scale. Molecular Biology and Evolution, 38(12), 5825–5829. 10.1093/molbev/msab293

Chaumeil, P.-A., Mussig, A. J., Hugenholtz, P., & Parks, D. H. (2022). GTDB-Tk v2: Memory friendly classification with the genome taxonomy database. Bioinformatics, 38(23), 5315–5316. 10.1093/bioinformatics/btac672

Chen, S. (2023). Ultrafast one-pass FASTQ data preprocessing, quality control, and deduplication using fastp. iMeta, 2(2), e107. 10.1002/imt2.107

Corso, D., Iburg, S., Coci, M., Cordone, A., Cesare, A. D., Giovannelli, D., Sbaffi, T., Martínez, A., & Eckert, E. M. (2026). Freshwater beach sandy sediments harbour latent β-lactamases with deep phylogenetic roots in Chitinophagaceae. Research Square. 10.21203/rs.3.rs-8368209/v1

Eddy, S. R. (2011). Accelerated Profile HMM Searches. PLOS Computational Biology, 7(10), e1002195. 10.1371/journal.pcbi.1002195

Eng, A., Verster, A. J., & Borenstein, E. (2020). MetaLAFFA: A flexible, end-to-end, distributed computing-compatible metagenomic functional annotation pipeline. BMC Bioinformatics, 21(1), 471. 10.1186/s12859-020-03815-9

Garfias-Gallegos, D., Zirión-Martínez, C., Bustos-Díaz, E. D., Arellano-Fernández, T. V., Lovaco-Flores, J. A., Espinosa-Jaime, A., Avelar-Rivas, J. A., & Sélem-Mójica, N. (2022). Metagenomics Bioinformatic Pipeline. In A. Pereira-Santana, S. D. Gamboa-Tuz, & L. C. Rodríguez-Zapata (Eds.), Plant Comparative Genomics (pp. 153–179). Springer US. 10.1007/978-1-0716-2429-6_10

Giovannelli, D. (2023). Trace metal availability and the evolution of biogeochemistry. Nature Reviews Earth & Environment, 4(9), 597–598. 10.1038/s43017-023-00477-y

Goussarov, G., Mysara, M., Vandamme, P., & Van Houdt, R. (2022). Introduction to the principles and methods underlying the recovery of metagenome-assembled genomes from metagenomic data. MicrobiologyOpen, 11(3), e1298. 10.1002/mbo3.1298

Gurevich, A., Saveliev, V., Vyahhi, N., & Tesler, G. (2013). QUAST: Quality assessment tool for genome assemblies. Bioinformatics, 29(8), 1072–1075. 10.1093/bioinformatics/btt086

Hera, M. R., Liu, S., Wei, W., Rodriguez, J. S., Ma, C., & Koslicki, D. (2024). Metagenomic functional profiling: To sketch or not to sketch? Bioinformatics, 40(Supplement_2), ii165–ii173. 10.1093/bioinformatics/btae397

Hyatt, D., Chen, G.-L., LoCascio, P. F., Land, M. L., Larimer, F. W., & Hauser, L. J. (2010). Prodigal: Prokaryotic gene recognition and translation initiation site identification. BMC Bioinformatics, 11(1), 119. 10.1186/1471-2105-11-119

Kang, D. D., Li, F., Kirton, E., Thomas, A., Egan, R., An, H., & Wang, Z. (2019). MetaBAT 2: An adaptive binning algorithm for robust and efficient genome reconstruction from metagenome assemblies. PeerJ, 7, e7359. 10.7717/peerj.7359

Krueger, F., James, F., Ewels, P., Afyounian, E., Weinstein, M., Schuster-Boeckler, B., Hulselmans, G., & Sclamons. (2023). *FelixKrueger/TrimGalore: V0.6.10 - add default decompression path* (Version 0.6.10) [Computer software]. [object Object]. 10.5281/ZENODO.7598955

Ladoukakis, E., Kolisis, F. N., & Chatziioannou, A. A. (2014). Integrative workflows for metagenomic analysis. Frontiers in Cell and Developmental Biology, 2. 10.3389/fcell.2014.00070

Langmead, B., Trapnell, C., Pop, M., & Salzberg, S. L. (2009). Ultrafast and memory-efficient alignment of short DNA sequences to the human genome. Genome Biology, 10(3), R25. 10.1186/gb-2009-10-3-r25

Li, D., Liu, C.-M., Luo, R., Sadakane, K., & Lam, T.-W. (2015). MEGAHIT: An ultra-fast single-node solution for large and complex metagenomics assembly via succinct de Bruijn graph. Bioinformatics, 31(10), 1674–1676. 10.1093/bioinformatics/btv033

Lloyd, K. G., Steen, A. D., Ladau, J., Yin, J., & Crosby, L. (2018). Phylogenetically Novel Uncultured Microbial Cells Dominate Earth Microbiomes. mSystems, 3(5), 10.1128/msystems.00055-18. https://doi.org/10.1128/msystems.00055-18

Menzel, P., Ng, K. L., & Krogh, A. (2016). Fast and sensitive taxonomic classification for metagenomics with Kaiju. Nature Communications, 7(1), 11257. 10.1038/ncomms11257

Meziti, A., Rodriguez-R, L. M., Hatt, J. K., Peña-Gonzalez, A., Levy, K., & Konstantinidis, K. T. (2021). The Reliability of Metagenome-Assembled Genomes (MAGs) in Representing Natural Populations: Insights from Comparing MAGs against Isolate Genomes Derived from the Same Fecal Sample. Applied and Environmental Microbiology, 87(6), e02593–20. 10.1128/AEM.02593-20

Mölder, F., Jablonski, K. P., Letcher, B., Hall, M. B., Tomkins-Tinch, C. H., Sochat, V., Forster, J., Lee, S., Twardziok, S. O., Kanitz, A., Wilm, A., Holtgrewe, M., Rahmann, S., Nahnsen, S., & Köster, J. (2021). Sustainable data analysis with Snakemake. F1000Research, 10, 33. 10.12688/f1000research.29032.1

Nelson, W. C., Tully, B. J., & Mobberley, J. M. (2020). Biases in genome reconstruction from metagenomic data. PeerJ, 8, e10119. 10.7717/peerj.10119

Nurk, S., Meleshko, D., Korobeynikov, A., & Pevzner, P. A. (2017). metaSPAdes: A new versatile metagenomic assembler. Genome Research, 27(5), 824–834. 10.1101/gr.213959.116

Pan, S., Zhu, C., Zhao, X.-M., & Coelho, L. P. (2022). A deep siamese neural network improves metagenome-assembled genomes in microbiome datasets across different environments. Nature Communications, 13(1), 2326. 10.1038/s41467-022-29843-y

Parks, D. H., Imelfort, M., Skennerton, C. T., Hugenholtz, P., & Tyson, G. W. (2015). CheckM: Assessing the quality of microbial genomes recovered from isolates, single cells, and metagenomes. Genome Research, 25(7), 1043–1055. 10.1101/gr.186072.114

Parks, D. H., Rinke, C., Chuvochina, M., Chaumeil, P.-A., Woodcroft, B. J., Evans, P. N., Hugenholtz, P., & Tyson, G. W. (2017). Recovery of nearly 8,000 metagenome-assembled genomes substantially expands the tree of life. Nature Microbiology, 2(11), 1533–1542. 10.1038/s41564-017-0012-7

Ricciardelli, A., De Pins, B., Brusca, J., Correggia, M., Di Iorio, L., Cascone, M., Giardina, M., Castaldi, S., Isticato, R., Iacono, R., Moracci, M., Nappi, N., Pollio, A., Vetriani, C., Leone, S., Cordone, A., & Giovannelli, D. (2025). *Trace metals availability controls terminal electron acceptor utilization in* Escherichia coli. Microbiology. 10.1101/2025.01.08.631794

Riesenfeld, C. S., Schloss, P. D., & Handelsman, J. (2004). Metagenomics: Genomic Analysis of Microbial Communities. Annual Review of Genetics, 38(1), 525–552. 10.1146/annurev.genet.38.072902.091216

Rogers, T. J., Buongiorno, J., Jessen, G. L., Schrenk, M. O., Fordyce, J. A., De Moor, J. M., Ramírez, C. J., Barry, P. H., Yücel, M., Selci, M., Cordone, A., Giovannelli, D., & Lloyd, K. G. (2023). Chemolithoautotroph distributions across the subsurface of a convergent margin. The ISME Journal, 17(1), 140–150. 10.1038/s41396-022-01331-7

Schwengers, O., Jelonek, L., Dieckmann, M. A., Beyvers, S., Blom, J., & Goesmann, A. (2021). Bakta: Rapid and standardized annotation of bacterial genomes via alignment-free sequence identification. Microbial Genomics, 7(11), 000685. 10.1099/mgen.0.000685

Sequeira, J. C., Rocha, M., Alves, M. M., & Salvador, A. F. (2022). UPIMAPI, reCOGnizer and KEGGCharter: Bioinformatics tools for functional annotation and visualization of (meta)-omics datasets. Computational and Structural Biotechnology Journal, 20, 1798–1810. 10.1016/j.csbj.2022.03.042

Shaffer, M., Borton, M. A., McGivern, B. B., Zayed, A. A., La Rosa, S. L., Solden, L. M., Liu, P., Narrowe, A. B., Rodríguez-Ramos, J., Bolduc, B., Gazitúa, M. C., Daly, R. A., Smith, G. J., Vik, D. R., Pope, P. B., Sullivan, M. B., Roux, S., & Wrighton, K. C. (2020). DRAM for distilling microbial metabolism to automate the curation of microbiome function. Nucleic Acids Research, 48(16), 8883–8900. 10.1093/nar/gkaa621

Sieber, C. M. K., Probst, A. J., Sharrar, A., Thomas, B. C., Hess, M., Tringe, S. G., & Banfield, J. F. (2018). Recovery of genomes from metagenomes via a dereplication, aggregation and scoring strategy. Nature Microbiology, 3(7), 836–843. 10.1038/s41564-018-0171-1

Tadrent, N., Dedeine, F., & Hervé, V. (2023). SnakeMAGs: A simple, efficient, flexible and scalable workflow to reconstruct prokaryotic genomes from metagenomes. F1000Research, 11, 1522. 10.12688/f1000research.128091.2

Tamames, J., & Puente-Sánchez, F. (2019). SqueezeMeta, A Highly Portable, Fully Automatic Metagenomic Analysis Pipeline. Frontiers in Microbiology, 9. 10.3389/fmicb.2018.03349

Tange, O. (2023). *GNU Parallel 20231122 ( Grindavík’)* [Computer software]. Zenodo. 10.5281/ZENODO.10199085

Tivey, M. (2007). Generation of Seafloor Hydrothermal Vent Fluids and Associated Mineral Deposits. Oceanography, 20(1), 50–65. 10.5670/oceanog.2007.80

Uritskiy, G. V., DiRuggiero, J., & Taylor, J. (2018). MetaWRAP—a flexible pipeline for genome-resolved metagenomic data analysis. Microbiome, 6(1), 158. 10.1186/s40168-018-0541-1

Van Damme, R., Hölzer, M., Viehweger, A., Müller, B., Bongcam-Rudloff, E., & Brandt, C. (2021). Metagenomics workflow for hybrid assembly, differential coverage binning, metatranscriptomics and pathway analysis (MUFFIN). PLOS Computational Biology, 17(2), e1008716. 10.1371/journal.pcbi.1008716

Wood, D. E., Lu, J., & Langmead, B. (2019). Improved metagenomic analysis with Kraken 2. Genome Biology, 20(1), 257. 10.1186/s13059-019-1891-0

Wu, Y.-W., Simmons, B. A., & Singer, S. W. (2016). MaxBin 2.0: An automated binning algorithm to recover genomes from multiple metagenomic datasets. Bioinformatics, 32(4), 605–607. 10.1093/bioinformatics/btv638

Yang, Y., Jiang, X., Chai, B., Ma, L., Li, B., Zhang, A., Cole, J. R., Tiedje, J. M., & Zhang, T. (2016). ARGs-OAP: Online analysis pipeline for antibiotic resistance genes detection from metagenomic data using an integrated structured ARG-database. Bioinformatics, 32(15), 2346–2351. 10.1093/bioinformatics/btw136

Yepes-García, J., & Falquet, L. (2024). Metagenome quality metrics and taxonomical annotation visualization through the integration of MAGFlow and BIgMAG. F1000Research, 13, 640. 10.12688/f1000research.152290.2

Yoo, A. B., Jette, M. A., & Grondona, M. (2003). SLURM: Simple Linux Utility for Resource Management. In D. Feitelson, L. Rudolph, & U. Schwiegelshohn (Eds.), Job Scheduling Strategies for Parallel Processing (pp. 44–60). Springer. 10.1007/10968987_3

Zhu, C., Miller, M., Marpaka, S., Vaysberg, P., Rühlemann, M. C., Wu, G., Heinsen, F.-A., Tempel, M., Zhao, L., Lieb, W., Franke, A., & Bromberg, Y. (2018). Functional sequencing read annotation for high precision microbiome analysis. Nucleic Acids Research, 46(4), e23. 10.1093/nar/gkx1209

